# Evolution and co-evolution of the suck behaviour, a postcopulatory female resistance trait that manipulates received ejaculate

**DOI:** 10.1101/2022.04.13.485945

**Authors:** Pragya Singh, Jeremias N. Brand, Lukas Schärer

## Abstract

Sexual conflicts over the post-mating fate of received ejaculate can favour traits in one sex that are costly to the other. Reciprocally mating hermaphrodites face unique challenges as they mate simultaneously in both the male and female role, potentially leading to receipt of unwanted ejaculate. Reciprocal mating can then give rise to postcopulatory female resistance traits that allow manipulation of received ejaculate. A putative example is the suck behaviour, observed in the flatworm genus *Macrostomum*. It involves the sperm recipient placing its pharynx over its own female genital opening and appearing to suck, likely removing received ejaculate after mating. The genus also contains hypodermically-inseminating species that presumably exhibit unilateral mating and have not been observed to suck. Here, we examine the evolution of the suck behaviour in *Macrostomum*, aiming to document the mating behaviour in 64 species. First, we provide videographic evidence that ejaculate is indeed removed during the suck behaviour in a reciprocally mating species, *Macrostomum hamatum*. Next, we show evolutionary positive correlations between the presence, duration and frequency of reciprocal mating behaviour and the suck behaviour, providing clear evidence that the suck behaviour co-evolves with reciprocal mating behaviour. Finally, we show an association between reproductive behaviour and reproductive morphology, suggesting that reproductive morphology can be used for inferring the behavioural mating strategy of a species. Together our study demonstrates sexual antagonistic coevolution leading to the evolution of a postcopulatory behavioural trait that functions as a female counter-adaptation allowing individuals to gain control over received ejaculate in a hermaphroditic sexual system.

## Introduction

Sexual conflict is defined as the conflict between the two sexes over their evolutionary interests involving reproduction (Charnov, 1979; Parker, 1979; Arnqvist & Rowe, 2005). The primordial cause of sexual conflict is anisogamy, in which the male sex produces more but smaller gametes (called sperm in animals), whereas the female sex produces fewer but larger gametes (called eggs in animals) (Parker, 2011). Because of this asymmetry, eggs are often a limiting resource for reproductive success, resulting in divergent interests between the two sexes (Bateman, 1948; Lehtonen *et al*., 2016). Furthermore, these conflicting interests can give rise to traits expressed by one sex that are costly to the other sex, resulting in antagonistic co-evolution between the sexes (Holland & Rice, 1998; Arnqvist & Rowe, 2005). Although work on sexual conflict has primarily focussed on separate-sexed organisms, sexual conflict is also pervasive in the lesser-studied hermaphroditic organisms (Charnov, 1979; Leonard, 1991; Michiels, 1998; Abbott, 2011; Schärer *et al*., 2015).

Of particular interest is the biology of simultaneous hermaphrodites (referred to as hermaphrodites hereafter), which involves unique sexual conflicts. For example, there can be conflicts between the mating partners over the sex role exhibited in a mating, namely mating as a sperm donor, a sperm recipient, or both. Depending on the costs and benefits of mating in each role, this may lead to sex role preferences (Michiels, 1998; Schärer *et al*., 2015). These are linked to Bateman’s principle, a term coined by Charnov (1979), which reflects the notion that there is a “greater dependence of males for their fertility on frequency of insemination” (Bateman, 1948). In his seminal paper, Charnov (1979) explored the proposal that Bateman’s principle also applies to simultaneous hermaphrodites. He concluded that, if true, hermaphroditic individuals may often mate more in order to give away sperm than to receive sperm, resulting in a mating conflict between the partners.

This conflict over the sex roles can be resolved via different mating strategies. One such strategy is reciprocal mating (also called reciprocal copulation), in which the partners simultaneously mate in both the male and female role. Each sperm donor is thus also a sperm recipient, and while multiple mating offers more opportunities to donate sperm, it may also lead to receipt of unwanted ejaculate from the partners. While this strategy seems like a cooperative conflict resolution, it could shift the conflict from the precopulatory to the postcopulatory arena (Schärer *et al*., 2015). In the presence of sperm competition, a donor—in order to secure a greater share of paternity—may often donate more sperm than the recipient requires for fertilisation, thereby potentially causing direct costs, such as a risk of polyspermy (Frank, 2000). But even if there are no direct costs posed by the received ejaculate, mating with multiple partners—which is probably the norm for most species (Jennions & Petrie, 2007; Kokko & Mappes, 2013; Arbuthnott *et al*., 2015)—could lead to the evolution of cryptic female choice (Charnov, 1979; Eberhard, 1996; Hemmings & Birkhead, 2017). Thus, receipt of excessive or unwanted ejaculate can favour the evolution of female resistance traits that allow postcopulatory control and rejection of the received ejaculate, e.g. via sperm digestion (Charnov, 1979).

Female resistance traits can in turn favour the evolution of male persistence traits, including other mating strategies. Such counter-adaptations may allow the sperm donor to either counteract or bypass the female resistance traits, thereby retain or regain access to the recipient’s eggs (Charnov, 1979; Schärer *et al*., 2015). An example of such an alternative mating strategy involves forced unilateral hypodermic insemination (also called hemocoelic insemination; Charnov 1979). Here one of the partners mates in the male role and donates sperm, while the other mates in the female role, potentially against its interests, and receives sperm hypodermically via a traumatic male copulatory organ (Lange *et al*., 2013; Reinhardt *et al*., 2015). With both types of mating strategy these sexual conflicts could then lead to the evolution of multiple male persistence and female resistance traits (spanning behaviour, morphology and physiology) that act jointly to either gain access to eggs, or to control and reject the received ejaculate, respectively (Arnqvist & Rowe, 2005). Therefore, we might expect behavioural mating strategies to be involved in sexually antagonistic coevolution, and thus to be correlated with morphological and/or physiological traits.

A putative example of a behavioural female resistance trait is the suck behaviour, originally documented in the free-living flatworm, *Macrostomum lignano* (Schärer *et al*., 2004). Studies in this reciprocally-mating simultaneous hermaphrodite have shown that matings are often followed by the suck behaviour, during which the worm bends down and places its pharynx over its own female genital opening (which is connected to the female antrum, the sperm-receiving organ) and then appears to suck. The suck behaviour is hypothesised to be a postcopulatory behaviour used for removing sperm or other ejaculate components received during mating and thus to function as a female resistance trait (Vizoso *et al*., 2010). However, while there have been multiple studies on this behaviour (Schärer *et al*., 2004, 2011, 2020; Marie-Orleach *et al*., 2013, 2017; Patlar *et al*., 2020), there has to date been no direct evidence for sperm and/or ejaculate actually being removed during the suck behaviour. Moreover, if the suck functions as a postcopulatory sexual selection process, it could affect the strength of sperm competition and potentially impact the optimal sex allocation (i.e. the amount of resources allocated to the male and female function) (van Velzen *et al*., 2009; Schärer & Pen, 2013). Indeed, studies have documented both inter- and intra-specific variation in sex allocation in *Macrostomum* (Singh *et al*., 2020b; Brand *et al*., 2022a; Singh & Schärer, 2021), with mating behaviour predicting the evolution of a species’ sex allocation (Brand *et al*., 2022a), but not the evolution of its sex allocation plasticity (Singh & Schärer, 2021).

Interestingly, *Macrostomum* species exhibit different combinations of reproductive morphological traits that are likely associated with the reciprocal mating and hypodermic insemination strategies (Figure 1A,B). Indeed, a previous study demonstrated an association between certain male and female reproductive traits and the mating strategy in 16 *Macrostomum* species, naming the two alternative outcomes the reciprocal and hypodermic mating syndrome, respectively (Schärer *et al*., 2011). A more recent study has used a refined composite measure, called the inferred mating syndrome, derived from the observation of additional components of the reproductive morphology, in an attempt to classify 145 *Macrostomum* species as showing either the reciprocal or hypodermic inferred mating syndrome, respectively (Figure 1B) (Brand *et al*., 2022b). The lateral bristles on the sperm in reciprocally mating species are hypothesized to represent a male persistence trait that allows the sperm to remain anchored in the female antrum and not be pulled out during the suck behaviour (Vizoso *et al*., 2010), whereas the thick female antrum wall might prevent internal injury resulting from the male genitalia during mating. In contrast, the sharp needle-like stylet tip of hypodermically inseminating species likely allows sperm injection through the partner’s epidermis, while the simple sperm design presumably aides its movement through the partner’s body (Schärer *et al*., 2011; Brand *et al*., 2022b).

**Figure 1.**
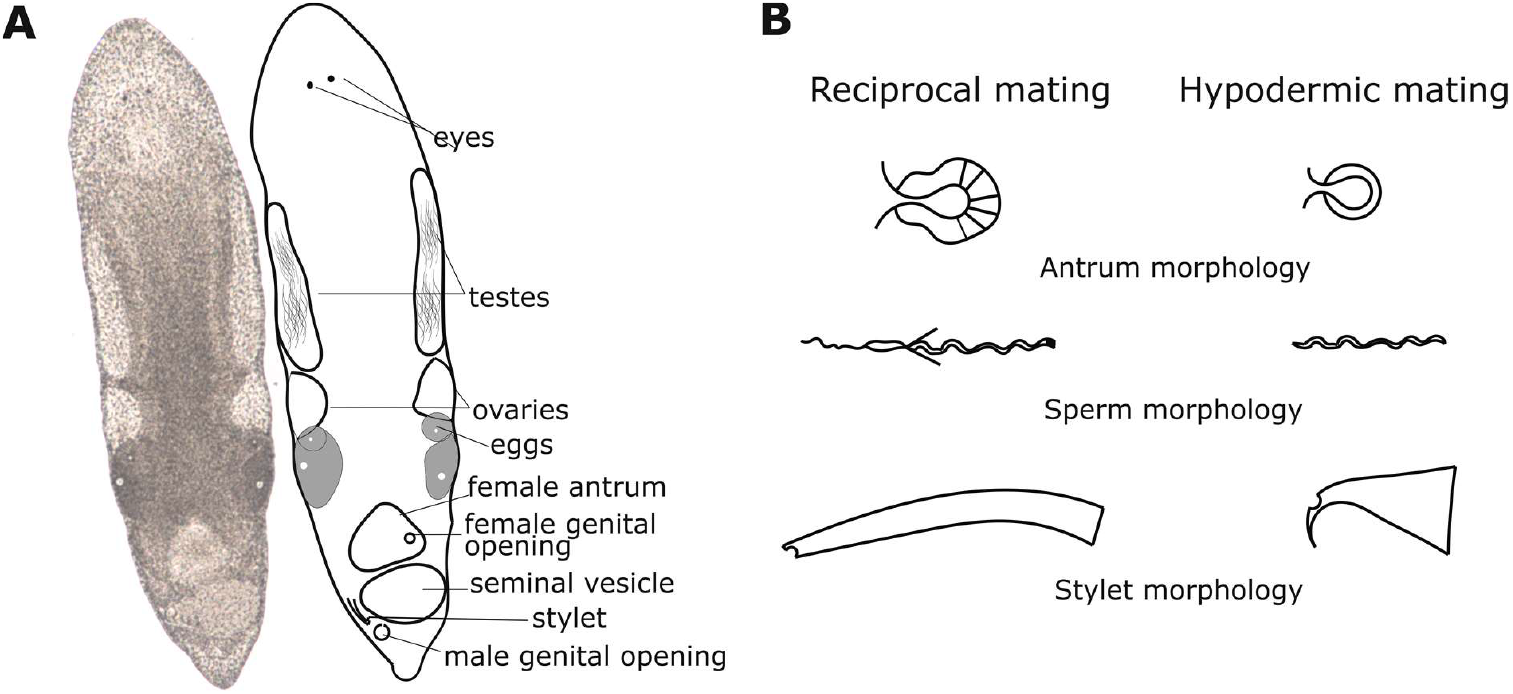
(A) Photograph and line drawing of an adult *Macrostomum cliftonense* (previously *M. cliftonensis*, name updated following Zhang *et al*., 2021), showing some of the components of the reproductive system to help understand the mating behaviour observations (total length ∼1.2 mm). (B) Schematic drawings of the typical morphology of the antrum (female reproductive organ), sperm, and stylet (male intromittent organ) of *Macrostomum* species with reciprocal mating (i.e. complex antrum and sperm, and stylet with a blunt distal end) and hypodermic mating (i.e. simple antrum and sperm, and a needle-like stylet) (see also Brand *et al*., 2022b).

Although sexual conflict has been studied in many organisms spanning different reproductive systems, studies on female resistance traits in hermaphrodites have been fewer, particularly in a phylogenetic context (Koene & Schulenburg, 2005; Beese *et al*., 2006, 2009; Anthes *et al*., 2008; Sauer & Hausdorf, 2009; Schärer *et al*., 2011; Brand *et al*., 2022b). In our study, we examine the evolution of the suck behaviour, aiming to document reproductive behaviour in a total of 64 *Macrostomum* species. As a result of this, we, for the first time, provide videographic evidence that ejaculate is indeed removed during the suck behaviour, supporting the previously proposed hypothesis for the function of this postcopulatory behaviour (Vizoso *et al*., 2010). Using this extensive behavioural data set, we examine correlations between different aspects of the mating and suck behaviour, and between reproductive morphology and the behavioural mating strategies, while accounting for the phylogenetic interrelationships. If the suck behaviour has indeed evolved as a postcopulatory strategy, we predict positive correlations between the presence, duration, and frequency of the mating behaviour and the suck behaviour. This could occur, e.g., if longer/frequent matings lead to more ejaculate being transferred, which would need longer/frequent sucks to remove the ejaculate. We might also expect a trade-off between copulation duration and frequency, if species that spend a lot of time in copulation cannot copulate that often, e.g. due to ejaculate limitation or mating taking up a lot of the total time (so that fewer mating could be done over a period, i.e., an autocorrelation). Similarly, if suck functions to remove ejaculate, we may expect a trade-off between suck duration and frequency, if shorter sucks necessitate the need for more frequent sucks to remove the ejaculate. Finally, we also expect the reproductive morphology to be a good proxy for inferring the behavioural mating strategy as a result of coevolution.

## Materials and Methods

### Study organisms

Species in the genus *Macrostomum* are small (∼0.3 to 3.0 mm body length) aquatic free-living flatworms that are highly transparent, permitting detailed observations of internal structures (for the general morphology see Figure 1A,B). The sperm and eggs are produced in the paired testes and paired ovaries, respectively, with studies documenting inter- and intra-specific variation in both testis and ovary size across the genus (Singh *et al*., 2020b; Brand *et al*., 2022a; Singh & Schärer, 2021). The female antrum is located anterior to the male antrum, connected to the outside, respectively, via a female genital opening (also female genital pore or vagina) and the male genital opening (also male genital pore). The stylet (male intromittent organ) resides within the male antrum and it is proximately connected via the vesicula granulorum (not shown) to the seminal vesicle, which contains sperm to be transferred during mating. In both reciprocally-mating and hypodermically-inseminating species, the female antrum serves as the egg-laying organ, while in reciprocally-mating species it additionally serves to receive the stylet during mating and as the sperm-storage organ (Vizoso *et al*., 2010; Schärer *et al*., 2011).

We obtained multiple specimens for a large number of *Macrostomum* species, collected from a range of locations and habitats, using a variety of extraction techniques, which we report on in more detail as part of separate studies on the phylogenetic interrelationships (Brand *et al*., 2022c) and reproductive character evolution in this genus (Brand *et al*., 2022b). Briefly, most specimens were sampled directly from natural field sites, while some were sampled from artificial ponds, or from aquaria containing other study organisms, and they were generally observed within a few days of collection. Other specimens were obtained from short- and long-term laboratory cultures maintained either by our group or by colleagues.

Following Brand *et al*. (2022b), 38 of a total of 64 species *Macrostomum* included in the current study were classified as exhibiting the reciprocal inferred mating syndrome, because they had a blunt tip of the stylet (the male intromittent organ), and of these all but one had received sperm in the antrum. A further 6 species with a sharp stylet were also classified as exhibiting the reciprocal inferred mating syndrome because they had complex sperm with lateral bristles and we observed sperm in the antrum (Figure 2). In contrast, 15 species were classified as exhibiting the hypodermic inferred mating syndrome because allosperm was exclusively found hypodermically. An additional 4 species without observation of received sperm were classified as exhibiting the hypodermic inferred mating syndrome because they had a simple female antrum (no thickening of the antrum wall and no visible cellular valve), a sperm design with reduced or absent bristles and a sharp stylet tip. Finally, one species (*Macrostomum sp*. 101) was classified as intermediate because received sperm was observed both in the antrum and within the tissue (Figure 2).

**Figure 2.**
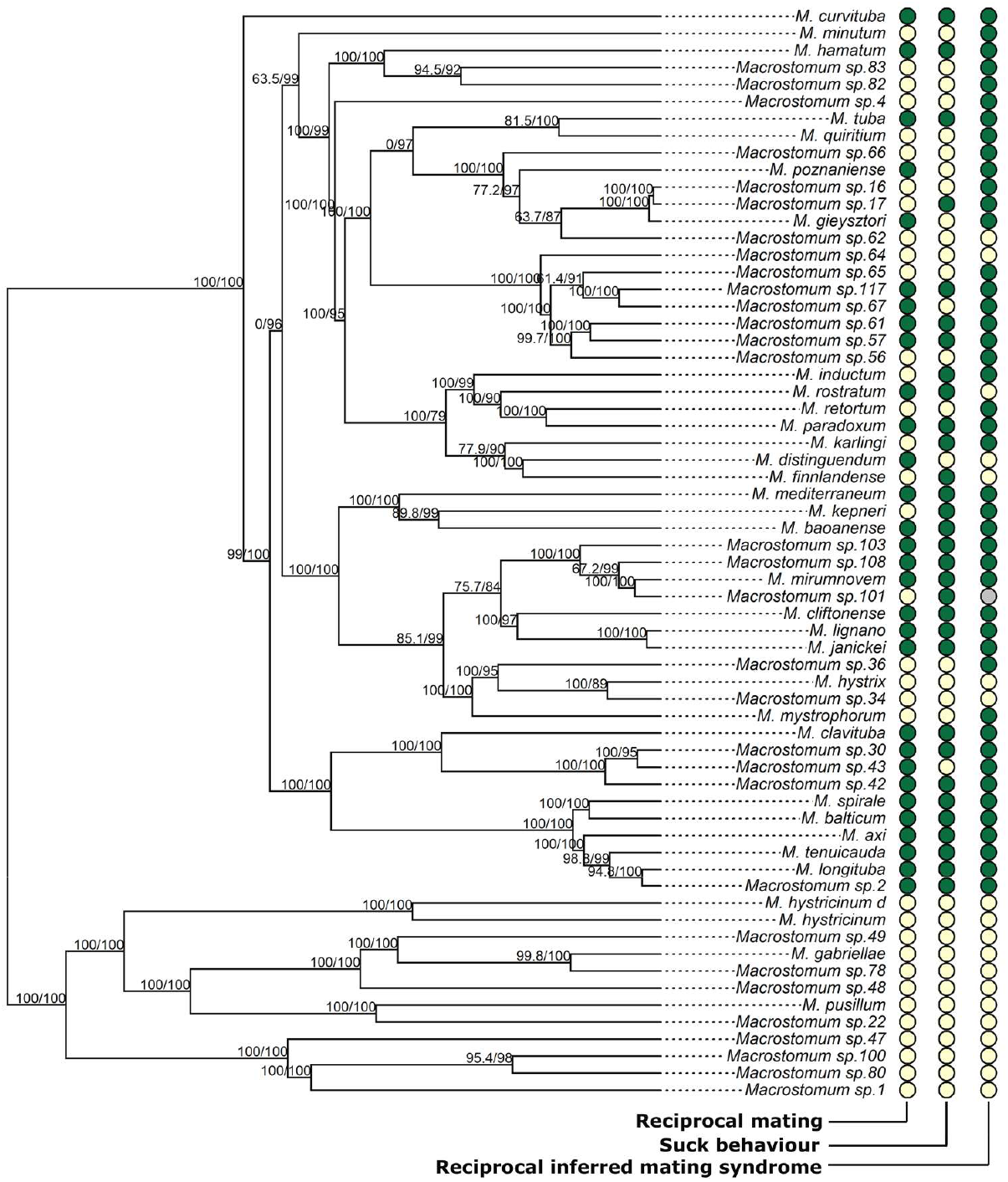
Presence (green) or absence (yellow) of reciprocal mating, the suck behaviour, and the reciprocal inferred mating syndrome across the *Macrostomum* phylogeny (for a total of 64 *Macrostomum* species, see Brand *et al*., 2022c for full phylogeny). Note that for the behaviourally-inferred traits an absence may be due to a lack of sufficient data for observing the behaviour, and that for the reciprocal inferred mating syndrome the absence represents the hypodermic inferred mating syndrome (except for *Macrostomum* sp. 101, which showed an intermediate inferred mating syndrome, grey). Branch supports are indicated by ultrafast bootstrap (first number) and approximate likelihood ratio tests (second number), respectively (from Brand *et al*., 2022c).

### Observation methodology

We aimed at documenting the mating behaviour of all 64 *Macrostomum* species, by placing the worms in mating chambers (Schärer *et al*., 2004). A mating chamber consisted of the worms being placed between two microscope slides in small drops (i.e. either freshwater or water with different salinity, depending on the collection habitat), with a certain number of spacers (separating the slides), and sealed with pure white Vaseline (note that we generally also placed 4-6 empty drops around, to reduce evaporation). We adjusted the spacer number and drop volume depending on the size and number of worms in a drop, respectively. Usually, for a pair of worms of the size of *M. lignano* (∼1.5 mm body length), we used 2 spacers (each spacer being ∼105 μm) and a drop size of ∼3 μl. Movies were recorded when specimens were available and therefore across several sampling campaigns. Consequently, the recording setups differed (macro lenses, cameras or lighting conditions). However from a previous detailed study of two *Macrostomum* species we know that these minor setup differences are unlikely to bias our observations (Singh *et al*., 2020a). Usually, the movies were recorded in QuickTime Format using BTV Pro (http://www.bensoftware.com/) at 1 frame s^-1^, but for some species we also generated detailed close-up movies, where worms were manually tracked at higher magnifications under a compound microscope and filmed at higher frame rates (see next section). All worms were visually checked for sexual maturity (defined as having visible gonads or eggs), either before or after filming.

### Detailed observation of mating and suck behaviour in Macrostomum hamatum

While earlier work documented sperm sticking out of the female antrum after the suck behaviour in *M. lignano* (Schärer *et al*., 2004, 2011), direct observations of ejaculate removal have not been reported to date. Here we could document ejaculate removal in detailed close-up movies of *M. hamatum*, possibly since field-collected specimens of this species appeared to be more transparent than other species. This allowed us to clearly visualise the deposition and subsequent removal of ejaculate during the mating and suck behaviour, respectively. Specifically, we examine the mating behaviour of *M. hamatum*, collected on 27. July 2017 directly in front of the Tvärminne Zoological Station, Finland (N 59.84452, E 23.24986), in a detailed close-up movie (Supplementary Movie S1). Note that while we describe and illustrate only one such instance (in an extract from a longer movie), we also observed ejaculate removal in other detailed close-up movies of *M. hamatum* that we also deposit (https://doi.org/10.5281/zenodo.6354683), and these observations corroborated our finding as described here.

### Scoring of mating and suck behaviour across species

We scored the mating behaviours from the mating movies by visual frame-by-frame analysis (Supplementary Table S1). A reciprocal mating was scored when the tail plates of two worms were in ventral contact and intertwined, such that the female antrum was accessible to the partner’s stylet and vice-versa, which would allow reciprocal transfer of ejaculate. In most species, the copulatory posture is accompanied by the pair being tightly interlinked (like two interlocking Gs, see Figure 3), and thus similar to the mating behaviour originally described for *M. lignano* (Schärer *et al*., 2004). Note that in some species the mating posture can deviate from that observed in *M. lignano* (see Supplementary Table S2), such as, for example, in *Macrostomum* sp. 57 and *Macrostomum* sp. 61 (Supplementary Movie S2A,B). The mating duration was measured from the frame when the tail plates were in ventral contact (and usually tightly intertwined), to the frame where the tail plates were no longer attached to each other. We defined behaviours as matings only if the pair was in the above-described posture for at least 3 s. The suck duration was measured starting from the frame when an individual placed its pharynx over its female genital opening, up to the frame where the pharynx disengaged. Note that in some cases, individuals do not lie on their side while sucking (as generally seen in *M. lignano*), which can sometimes make it more difficult to observe the suck behaviour. For each replicate drop, we divided the total number of matings and sucks by the number of worms and the movie duration to obtain a standardized value. We then averaged the frequency and duration estimates across all replicate drops for each species to obtain the species estimates of the respective behaviours.

**Figure 3.**
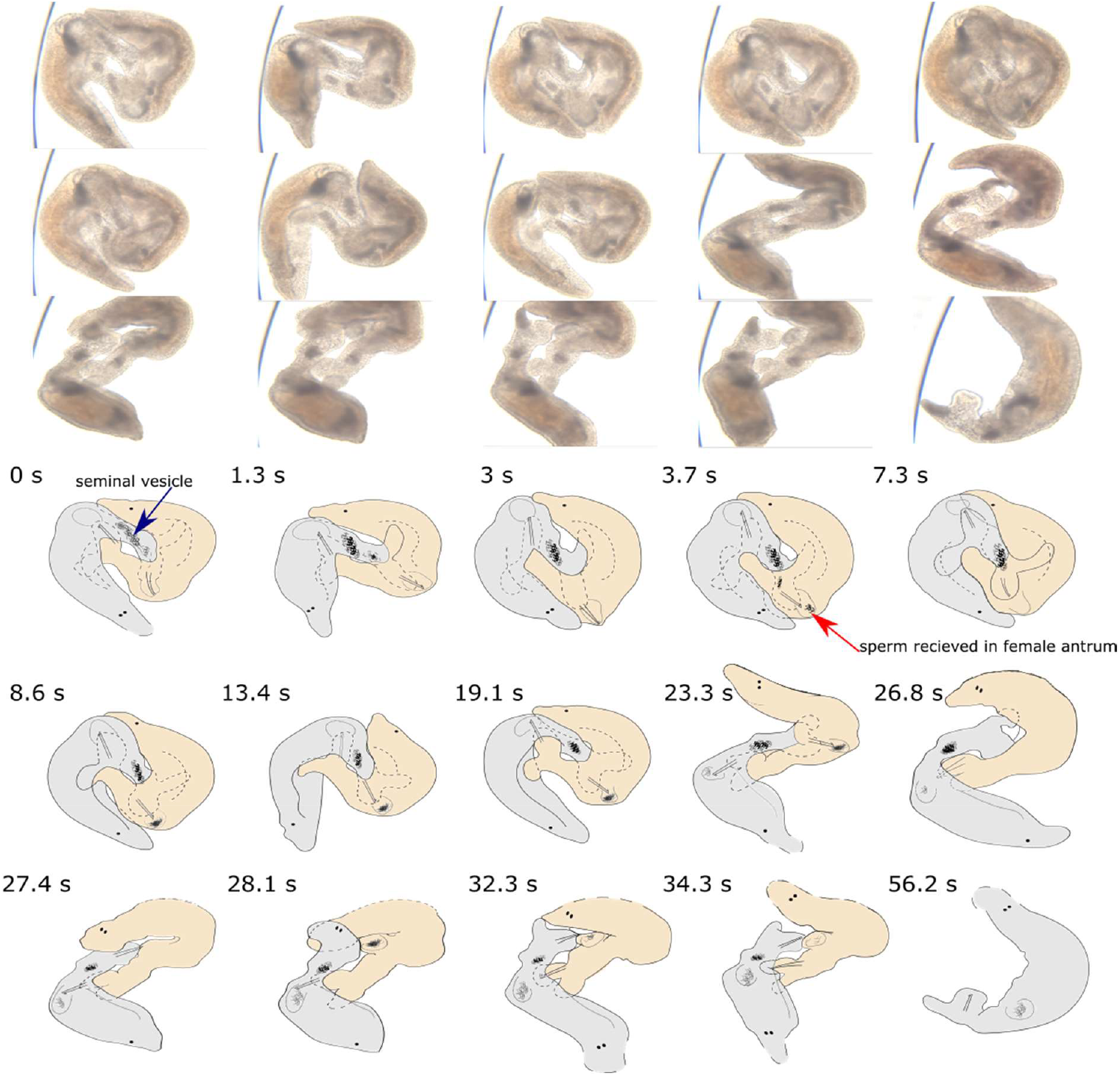
Reciprocal mating followed by a postcopulatory suck in *Macrostomum hamatum*, including ejaculate deposition by Grey (the worm on the left at 0 s) and its subsequent removal during the suck behaviour by Orange (the worm on the right at 0 s). Before transfer, the sperm is stored in the seminal vesicle of Grey (blue arrow in first frame), which is connected to its stylet. Ejaculate (dark mass indicated by red arrow) can be seen being deposited by Grey from the seminal vesicle starting from 3.7 s in the female antrum of Orange, followed by Orange pushing its female antrum region out (at 27.4 s) and sucking (note that Orange is also depositing ejaculate in Grey from 23.3 s). There is a visible reduction in the quantity of received ejaculate in the female antrum of Orange after the suck ends. Note that we call the frame from where we start describing the movie as t = 0 s, but the mating had already started before that timepoint. In some frames, parts of the worms are not visible on the video, and the presumed outlines are drawn using stippled lines. A high-resolution version is provided in Supplementary Figure S1A.

While we also invested significant effort into observing hypodermic insemination (see Table S1 and Results), we only saw some rare behavioural instances in a few species that could possibly represent cases of hypodermic insemination, such as, for example, in *Macrostomum* sp. 1 and *M. gabriellae* (Supplementary Movie S3A,B). Possible reasons for not observing hypodermic insemination could be that in many species such matings occur very rapidly or that they mate less frequently, possibly since they try to avoid sperm receipt (Apelt, 1969; Michiels, 1998). Given that we could not confirm the presence of hypodermic insemination, we scored species either as having reciprocal mating being present (when it was observed) or absent (when it was not observed) (Figure 2). Note, however, that the absence of observations of reciprocal mating does not necessarily imply the presence of hypodermic insemination. Instead, it could also result from a reciprocally-mating species not mating under laboratory conditions and/or from an overall low mating frequency of a species. Similarly, while the presence of the suck behaviour can clearly be identified in many species, the absence of observations of the suck behaviour does not necessarily mean that a species never shows this behaviour.

### Evolution of the mating and suck behaviour across the genus Macrostomum

To perform phylogenetic comparative analyses, we used a trimmed version of a recently published ultrametric large-scale phylogeny of the genus *Macrostomum* (i.e. the C-IQ-TREE phylogeny of Brand *et al*., 2022c). This phylogeny is based on an amino acid alignment of 385 genes from 98 species, supplemented with Sanger sequences from a *28S rRNA* fragment, which allowed the addition of a further 47 species, and calculated using a maximum likelihood approach (Brand *et al*., 2022c), covering all the species we included in the current study. Specifically, we determined 1) whether the presence/absence of reciprocal mating is correlated with the presence/absence of the suck behaviour, 2) whether the presence/absence of reciprocal mating is correlated to the presence/absence of the reciprocal inferred mating syndrome, and 3) whether there are correlations between the frequency and the duration of the reciprocal mating and suck behaviours among the species that show these behaviours.

#### Presence/absence of reciprocal mating and the suck behaviour

We used the DISCRETE model in BayesTraits V.3.0.1 to test for correlated evolution between reciprocal mating and the suck behaviour (both scored as present/absent), using the Reversible Jump Markov Chain Monte Carlo (RJ MCMC) approach (Pagel, 1994; Pagel & Meade, 2006; Meade & Pagel, 2016). Specifically, we compared the marginal likelihood of a dependent model, in which the presence of suck depends on the presence of reciprocal mating, to an independent model, in which the suck behaviour and reciprocal mating evolve independently. Each RJ MCMC chain was run for twelve million iterations and the first one million iterations were discarded as burn-in, after which the chain was sampled every 1000^th^ iteration. We used a gamma hyperprior (gamma 0 1 0 1), and placed 1000 stepping stones (with each iterating 10000 times) to obtain the marginal likelihood values for the models. We ran three separate chains each for the dependent and independent model to check for the stability of the likelihood values and convergence. Using the R package coda (Plummer *et al*., 2006), we confirmed that the chains had converged (Gelman & Rubin, 1992; Brooks & Gelman, 1998) and that the Effective Sample Size was >200 for all parameters. In addition, we also confirmed that the acceptance rate was between 20-40% (Pagel & Meade, 2006). We compared the alternative models with the Log Bayes Factor (BF), using the convention that BF values > 2 are considered as positive support for the best-fit model, while values between 5-10 and > 10 are considered as strong and very strong support for the model, respectively (Pagel & Meade, 2006). To examine the robustness, we repeated the analysis for a reduced dataset, by excluding six species that had in total been observed for < 21 h (∼10% quantile). For the dependent models of the full dataset, we estimated the transition rates among the different trait states by calculating Z values. This value can be understood as the percentage of times a transition rate was set to zero, with a high value thus indicating that the transition between two states is unlikely. We expect a correlation between the presence of reciprocal mating and the presence of the suck behaviour, which would corroborate that the suck behaviour indeed is a postcopulatory behaviour that is linked to reciprocal mating, rather than possibly serving a function that is also present in species with hypodermic insemination.

#### Presence/absence of reciprocal mating and the reciprocal inferred mating syndrome

We checked for an association between reciprocal mating (scored as present/absent) and the inferred mating syndrome (scored as reciprocal/hypodermic), using the DISCRETE model in BayesTraits V.3.0.1 (as above). One of the species, *Macrostomum* sp. 101, had a morphology that was scored intermediate between reciprocal and hypodermic (Brand *et al*., 2022b), but since the discrete method in BayesTraits only allows binary trait states, we excluded this species from this analysis. We expect a correlation between the presence of reciprocal mating behaviour and the reciprocal inferred mating syndrome, which could indicate that behaviour and morphology coevolve.

#### Correlations between the frequency and the duration of mating behaviours

In preparation for phylogenetic correlation analyses we estimated the phylogenetic signal for the continuous traits (i.e. the duration and frequency of both the reciprocal mating and the suck behaviour; log-transformed for all the analyses) using Pagel’s λ (Pagel, 1999; Revell, 2012). A λ value of 1 indicates a strong phylogenetic signal, while a value around 0 indicates no/low phylogenetic signal (Pagel, 1999). We found phylogenetic signal that was significantly different from 0 for the suck frequency (λ=0.67, P=0.02), the suck duration (λ=0.76, P=0.005), and the reciprocal mating frequency (λ=0.50, P=0.05), but only marginally so for the mating duration (λ=0.46, P=0.06). For each trait, we then fitted four different models of trait evolution, i.e. Brownian motion, Ornstein–Uhlenbeck, Early-burst, and Lambda models (Harmon *et al*., 2008). We found that the Lambda model had the highest sample-size corrected Akaike Information Criterion (AICc) weights (Φι) (Supplementary Table S3), and this model was hence chosen for further PGLS analysis.

For the species that exhibited both reciprocal mating and the suck behaviour, we then investigated if there was a correlation between the frequency and duration of the reciprocal mating and suck behaviours, using phylogenetic generalized least-squares (PGLS) regression implemented in the *caper* package version 1.0.1 (Orme *et al*., 2014). PGLS accounts for the non-independence of the data by incorporating the phylogenetic relationships between species into the error structure of the model. For each analysis using the frequency and duration of the reciprocal mating and suck behaviours, the phylogenetic signal (Pagel’s λ) was estimated using the maximum likelihood approach. We examined the residuals of each model for normality and homogeneity (Mundry, 2014). Additionally, we scrutinized for influential cases (species) in each PGLS model, by excluding one species at a time from the data and rerunning the analysis, and comparing the results obtained with the results for the entire dataset (Mundry, 2014). And finally, we evaluated the robustness of our results by repeating the PGLS for a reduced dataset, which excluded five species in which mating or suck had only been observed in one replicate (note that this reduced dataset is different from the reduced dataset used in the above BayesTraits analysis).

We performed our analysis in R, version 3.6.1 (R Core Team, 2019).

## Results

### Sperm deposition and removal during mating and suck behaviour in Macrostomum hamatum

The general anatomy of the reproductive organs of *M. hamatum* is similar to that of many other reciprocally-mating *Macrostomum* species (Figure 1A,B). In the detailed movie of *M. hamatum*, the worms are already interlinked in the reciprocal copulatory position at the beginning of the clip (Figure 3, Supplementary Movie S1), and we consider this as t = 0 s (hereafter we refer to the worm on the right as Orange and the worm on the left as Grey, respectively). At this timepoint “the tail plates touch each other ventrally in opposing directions, while the anterior ventral surface of each worm touches the posterior dorsal surface of the partner”, as previously described for the copulatory position in *M. lignano* (Schärer *et al*., 2004). Interestingly, in *M. hamatum* the copulatory position resembles a square with rounded corners, as opposed to *M. lignano*, where it is more circular. This may in part be due to a strikingly different position of the tail plate, which in *M. hamatum* stands at a 90° angle from the posterior body axis and appears to poke into the anterior ventral surface of the partner, leading to a dorsal bulge in both Orange and Grey.

Moreover, *M. hamatum* has a much more prominent erection (i.e., a translucent finger-like structure on the ventral tail plate, likely formed by the eversion of the muscular male antrum), which pokes into the posterior ventral surface of the partner in the region of the female genital opening (although it is unclear if the erection actually enters the partner). The stylet of Grey— while moving inside of the relatively stationary erection—then performs poking movements that are directed towards Orange’s female antrum, initially without any transfer of ejaculate. At t = 3-5 s, the stylet of Grey is seen repeatedly poking against the dorsal side of Orange’s female antrum wall, each time leading to a visible bulge on Orange’s dorsal side. In some of these frames one can see the sharp hook-shaped distal end of the stylet that is typical for *M. hamatum*. Eventually, Grey begins to deposit ejaculate (seen as a visible darkening of Orange’s female antrum lumen starting at about t = 5s). During this process the seminal vesicle of Grey empties (as seen at the base of the erection, see also drawing in Figure 3 at 0 s for location), while the female antrum of Orange fills up with ejaculate over the next ∼21 s (see Figure 3 from 3.7 s). Note that we here mainly focus on the sperm transfer from Grey to Orange, but in the meantime, Orange also pokes and eventually enters the female antrum of Grey (t = 16-20 s) and sperm is also transferred from Orange to Grey (between t = 21-27 s), although this is more difficult to follow in the movie.

At t = 28 s, Orange pushes out its female antrum region, places its pharynx over its female genital opening, and then sucks. The received ejaculate can be seen leaving Orange’s female antrum (i.e. the visible darkening in the female antrum lumen moves towards the pharynx between t = 29-30s). In total, the suck behaviour lasts for 7 s. Interestingly, during the suck the stylet of Grey remains anchored in Orange’s female genital opening (probably involving the above-mentioned hook). At t = 52 s, the mating ends after a mating duration of ∼64 s (recall that the worms were already in copula at t = 0 s). At t = 56 s, only Grey is in frame and the received ejaculate in its female antrum is clearly visible. It continues to have a small erection despite the mating being over. At t = 78 s, Grey pushes its female antrum region out and some sperm is ejected from the female antrum at t = 80 s, notably before the pharynx makes contact (Figure 4, Supplementary Movie S1). At t = 81 s, Grey puts its pharynx over its female genital opening and then sucks for 10 s. After the suck, some sperm can still be seen sticking out of the female antrum (similar to *M. lignano*, Schärer *et al*., 2004), especially at 92 s, but most of the ejaculate has been removed from the female antrum. The female antrum remains slightly everted and the erection somewhat visible until at least 108 s.

**Figure 4.**
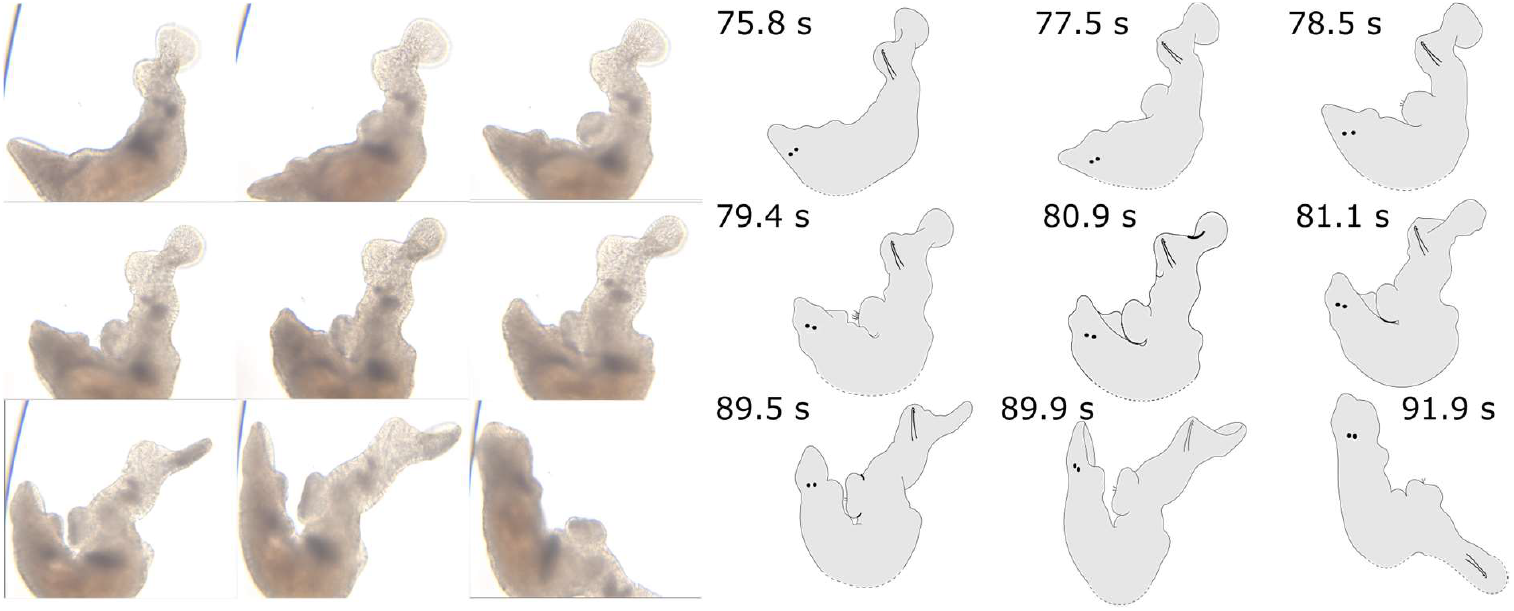
A postcopulatory suck following the reciprocal mating shown in Figure 3 (a continuation of the same movie), performed by Grey (i.e. the individual that had not yet sucked). Grey completely pushes out its female antrum region at t = 78 s (which leads to some sperm appearing near the female genital opening at t = 78.5 s), puts its pharynx over the female genital opening, and then sucks out most of the previously deposited ejaculate over a period of 10 s (from t = 81 s). Moreover, some sperm can be seen sticking out of the female genital opening after the suck ends. In some frames, part of the worm is not visible on the video, and the presumed outline is thus drawn using stippled lines. A high-resolution version is provided in Supplementary Figure S1B.

### Evolution of the mating and suck behaviour across the genus Macrostomum

We observed a total of 2796 worms across 64 *Macrostomum* species, with a mean of 44 worms and 76.7 hours of observation time per species, for a total observation time of 4908 hours. Of the 64 species, 30 species exhibited reciprocal mating behaviour, 31 species exhibited the suck behaviour, and 25 species exhibited both the reciprocal mating and suck behaviour (Figure 2).

#### Presence/absence of reciprocal mating and the suck behaviour

We found very strong support for the dependent model over the independent model of evolution for the correlation between the presence of reciprocal mating and the presence of the suck behaviour, with all three runs for each model providing highly consistent values (average marginal likelihood, independent=-89.25, dependent=-83.37, BF: 11.75; see also Supplementary Table S4a). This showed that the presence of reciprocal mating and the presence of the suck behaviour are strongly correlated. And the result was robust to observation time, since excluding the 6 species that were observed for < 21 h gave similar results (Supplementary Table S4a).

The transitions from the absence of both the reciprocal mating and suck behaviour to the presence of either of these traits were found to be the most unlikely, as is evident from the low transition rates and the high Z values (Figure 5a). Interestingly, the other transitions, including losing reciprocal mating or the suck behaviour from the state when they are both present, are all similarly likely. This contrast suggests that once both reciprocal mating and suck are lost or absent in a species, it is highly unlikely to regain either.

**Figure 5.**
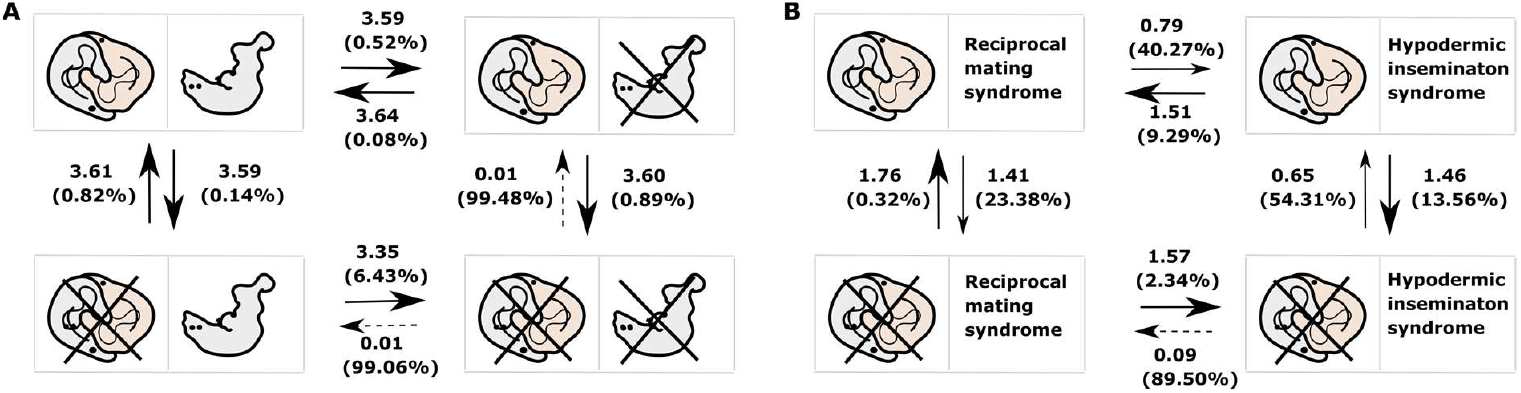
Correlated evolution of behavioural character states. The panels show the transition rates and the Z values (in brackets, expressed as %) for transitions between (A) the presence or absence of reciprocal mating and the suck behaviour (crossed out when absent), and (B) the presence or absence of reciprocal mating (crossed out when absent) and the inferred mating syndrome (from Brand *et al*., 2022c). For the transition rates, the mean of the posterior distributions across all runs is given. The Z value can be understood as the percentage of times the transition rate was set to zero, amongst all the sampled parameters. The different arrows represent different probabilities of transitions between the states: high probability (strong black arrows, Z value < 15%), moderate probability (thin black arrows, Z value 20-55%), and low probability (dashed black arrows, Z value > 85%). The posterior distributions of the transition rate parameters are given in Supplementary Figure S2.

#### Presence/absence of reciprocal mating and the reciprocal inferred mating syndrome

There was a clear correlation between the presence of reciprocal mating behaviour and the reciprocal inferred mating syndrome, as evident from the strong support for the dependent model over the independent model of evolution, with similar values for the three independent runs of each model (average marginal likelihood, independent=-69.09, dependent=-65.60, BF: 6.99; see also Supplementary Table S4b), suggesting that the reproductive morphology of a species can serve as a good proxy for its mating behaviour. And as before, our result was robust, as the reduced dataset gave us similar results (Supplementary Table S4b).

Transitions from the presence of reciprocal mating and the presence of the reciprocal inferred mating syndrome to the absence of either were moderately likely, while the converse transitions were very likely (Figure 5b). Similarly, transitions from the absence of reciprocal mating and the absence of the reciprocal inferred mating syndrome to the presence of either were either unlikely or relatively unlikely, while the converse transitions were very likely. Together this suggests that there is a strong association between reciprocal mating behaviour and morphological traits characterizing the reciprocal inferred mating syndrome (and between absence of reciprocal mating behaviour and the hypodermic inferred mating syndrome), such that species are attracted to these states and evolve away from states where the morphology and behaviour are mismatched.

#### Correlations between the frequency and the duration of mating behaviours

Among the species that exhibited reciprocal mating (n=30), the average mating frequency was 0.84 hr^-1^ (range: 0.02-7.82 hr^-1^, Figure 6A) and the average mating duration was 283.7 s (range: 5.2-4609 s, Figure 6B), with some sibling species showing fairly divergent values. Moreover, among the species that showed the suck behaviour (n=31), the average suck frequency was 0.54 hr^-1^ (range: 0.01-3.7 hr^-1^, Figure 6A) and the average suck duration was 9.6 s (range: 4.7-16.1 s, Figure 6C).

**Figure 6.**
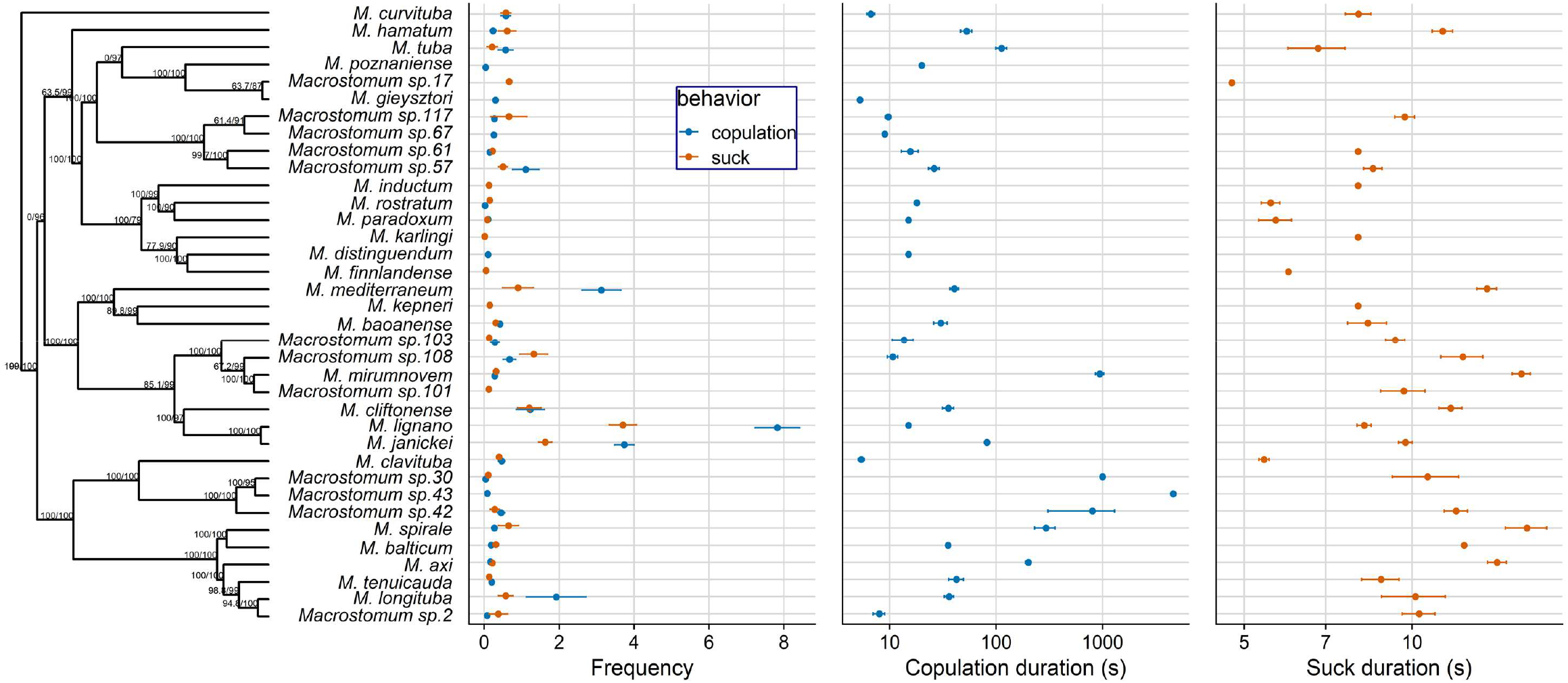
Trimmed phylogeny of the 36 *Macrostomum* species that showed reciprocal mating and/or the suck behaviour alongside data on means and standard errors of (A) reciprocal mating and suck frequency, (B) reciprocal mating duration (log-transformed), and (C) suck duration (log-transformed). Note that some species exhibited either only reciprocal mating or only the suck behaviour. Also note that for the species in which a behaviour had been observed in only 1 replicate, we report only that single value. The branch support values are indicated by ultrafast bootstrap (first number) and approximate likelihood ratio tests (second number), respectively (from Brand *et al*., 2022c).

In line with our predictions, we found significant positive relationships between both reciprocal mating frequency and suck frequency (Figure 7A), and reciprocal mating duration and suck duration (Figure 7B); while there was no significant relationship between reciprocal mating frequency and reciprocal mating duration (Figure 7C), and suck frequency and suck duration (Figure 7D). The reduced dataset also gave qualitatively similar results for all analysis (Supplementary Table S5).

**Figure 7.**
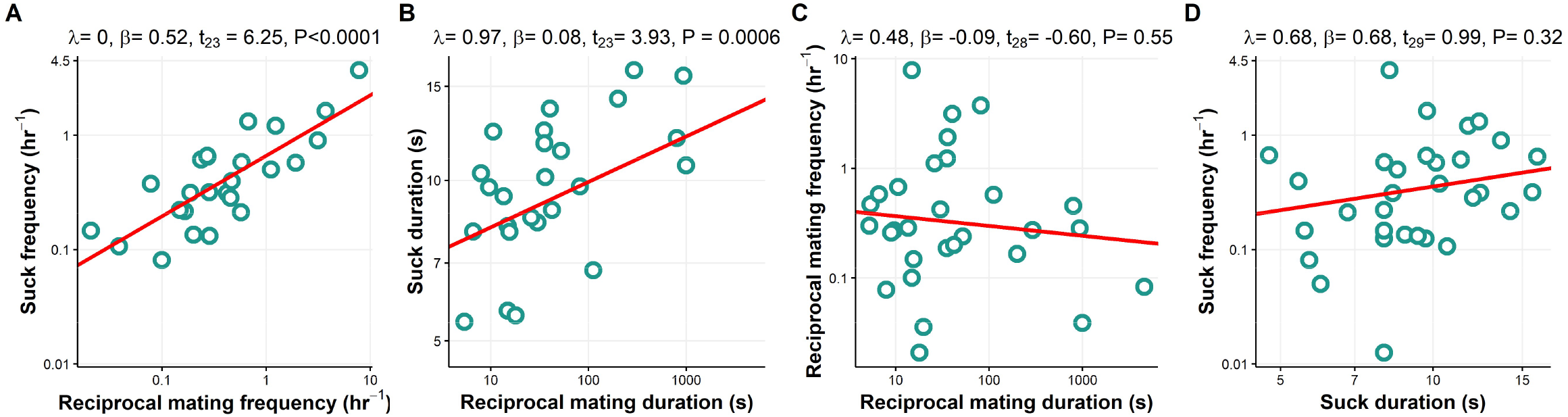
Relationships between (A) reciprocal mating frequency and suck frequency, (B) reciprocal mating duration and suck duration, (C) reciprocal mating duration and frequency, and (D) suck duration and frequency for *Macrostomum* species. Note that (A-D) show values plotted on log-transformed axes with PGLS results.

## Discussion

Sexual conflict can give rise to antagonistic coevolution in all sexual systems (Charnov, 1979; Bedhomme *et al*., 2009). Here we documented the widespread occurrence of a putative female resistance trait, the suck behaviour, in >30 species in the hermaphroditic flatworm genus *Macrostomum*. Moreover, the direct observation of ejaculate removal in one species, *M. hamatum*, corroborates the hypothesis that the suck functions as a female resistance trait to remove received ejaculate (Schärer et al. 2004; Vizoso et al. 2010; Schärer et al. 2011), and this interpretation is also supported by significant evolutionary correlations between different aspects of reciprocal mating and suck behaviour. Finally, we could also show that the reproductive morphology is a good proxy for inferring the mating strategy of a species, presumably also as a result of coevolution. In the following we discuss these findings in more detail.

### Sperm deposition and removal during mating and suck behaviour in Macrostomum hamatum

While multiple studies in *Macrostomum* have examined aspects of the suck behaviour (Schärer *et al*., 2004, 2011, 2020; Marie-Orleach *et al*., 2013, 2017; Patlar *et al*., 2020; Singh *et al*., 2020a), its involvement in removing received ejaculate components has so far only been hypothesized. Our detailed observations of mating interactions in *M. hamatum* provide the first direct evidence that ejaculate is indeed removed during this postcopulatory behaviour. Interestingly, compared to *M. lignano* (Schärer *et al*., 2004), *M. hamatum* has a more rectangular mating posture (possibly due to the angular position of the tail plate), a larger erection around the stylet, and the worms prominently evert the female antrum just before the suck behaviour, likely as a result of muscular contractions. This could result from differences in the female antrum morphology: while *M. hamatum* has a strong musculature and an inner second chamber connecting to the main female antrum (Luther, 1947), *M. lignano* has a somewhat simpler female antrum with a single chamber (Ladurner *et al*., 2005; Vizoso *et al*., 2010). Similarly, the prominent erection of the male antrum could result from a muscular morphology that is similar to the muscular cirrus seen in species of the sister genus, *Psammomacrostomum* (Ax, 1966; Janssen *et al*., 2015). The combination of a rather prominent female antrum and the relatively transparent specimens may have helped us visualise the function of the suck behaviour better in *M. hamatum* than in other *Macrostomum* species observed to date.

While we see ejaculate being removed during the suck behaviour, we cannot clearly determine whether it is ingested. Although sperm digestion is widespread in hermaphrodites (Charnov, 1979; Baur, 1998; Dillen *et al*., 2009; Koene *et al*., 2009), it usually occurs inside an organ connected to the individual’s reproductive system, unlike in the case of the suck behaviour. To our knowledge, there have been only two earlier reports of sperm being orally taken up in hermaphrodites, one in the arrow worm *Spadella cephaloptera* (John, 1933) and the other in the leech *Placobdella parasitica* (Myers, 1935). Thus, the suck behaviour seems to be a novel trait, which to date has only been observed in species of the Macrostomidae (including a member of the sister genus *Psammomacrostomum*; P. Singh, pers. obs.). Similar to the suck behaviour, females of the ladybird beetle, *Adalia bipunctata*, consume a spermatophore after mating (Perry & Rowe, 2008). Moreover, there is also sperm dumping in many separate-sexed species, in which the female physically ejects received sperm from her reproductive tract, and this, at least in some cases, is thought to be a mechanism of cryptic female choice (Snook & Hosken, 2004; Peretti & Eberhard, 2010; Firman *et al*., 2017). If the suck behaviour also functioned in cryptic female choice, we might expect individuals to remove or retain sperm of certain partners more frequently (e.g. Pizzari & Birkhead, 2000). This has also been observed in *M. lignano*, where the propensity of the recipient to suck is affected by the mating status (Marie-Orleach *et al*., 2013) and the genotype of its partners (Marie-Orleach *et al*., 2017). However, it is difficult to ascertain whether the suck implies an active choice by the recipient or whether it is sometimes prevented as a result of a manipulation by the donor (Patlar *et al*., 2020). Moreover, our study documents in detail the reciprocal transfer and deposition of sperm by both mating individuals during a reciprocal mating in *Macrostomum* (but see Ax & Borkott, 1968 which documents mating and unilateral sperm transfer in *M. salinum*, now considered to be *M. romanicum*).

### Evolution of the mating and suck behaviour across the genus Macrostomum

We found a significant evolutionary correlation between the presence of reciprocal mating and the suck behaviour (Figure 5a). In reciprocally-mating species, ejaculate is deposited in the female antrum allowing its removal during the suck behaviour, while in hypodermically-inseminating species, sperm is injected potentially anywhere in the body (Schärer *et al*., 2011; Brand *et al*., 2022b). Given that the function of the suck behaviour indeed appears to be the removal of ejaculate, we do not expect to see the suck behaviour in hypodermically-inseminating species. Performing a suck at a site of hypodermic insemination might not permit effective ejaculate removal (particularly also given the above-mentioned active participation of the female antrum musculature), but instead would more likely lead to additional tissue damage. Interestingly, the transition rates showed that while it is unlikely for a species that lacks both the reciprocal mating and suck behaviour to gain either of these traits, the loss of either reciprocal mating or the suck behaviour was estimated as being more likely. These transitions could represent transitional steps towards hypodermic insemination, which might arise as a means to bypass the female control and allow access to the eggs (Charnov, 1979; Brand *et al*., 2022b). Moreover, this interpretation is also supported by the finding that there are multiple origins of hypodermic insemination in the genus *Macrostomum* (Brand *et al*., 2022b; Singh & Schärer, 2021). There are at least nine independent shifts from reciprocal mating to hypodermic insemination in *Macrostomum*, while no transition is observed in the converse direction (Brand *et al*., 2022b).

However, it is important to point out that some of these findings could also have resulted from a lack of observations of either the reciprocal mating and/or suck behaviour (despite being present in a species), leading to an overestimation of these transition rates. Specifically, there were six species that showed only the reciprocal mating behaviour and five species that showed only the suck behaviour (Supplementary Table S1). These mismatches usually appeared in species for which we had comparatively few observation hours (for more detail see Supplementary Figure S3), suggesting that additional observations could help to further ascertain the actual presence/absence of reciprocal mating or the suck behaviour, respectively. Moreover, mismatches could result from a species not exhibiting some behaviours under our laboratory conditions, or they might indicate that a species indeed lacks a behaviour. In addition, if a species mates only rarely, individuals might be less inclined to remove the sperm they receive, and in our study the species that showed reciprocal mating but did not suck, had low or intermediate mating frequencies (see *Macrostomum* sp. 43, *Macrostomum* sp. 67, *M. distinguendum, M. gieysztori*, and *M. poznaniense* in Figure 6A). Alternatively, species might actually lack reciprocal mating, but losing a resistance trait like the suck behaviour might take longer, particularly if the suck behaviour does not impose costs on the fecundity. Moreover, the suck behaviour could have additional functions, such as possibly removing egg material that remains in the antrum after egg laying. Species are predicted to lose defensive or resistance traits only after the persistence traits have become substantially less harmful, leading to a time lag (Parker, 1979). A study on the seed beetle, *Callosobruchus maculatus*, showed that, while large males evolved relatively reduced length of genital spines under monogamy, there was no detectable evolution in female genitalia within the same time period (Cayetano *et al*., 2011). And finally, since the worms we observed may often have mated before we placed them into the mating chambers, some of the observed sucks might have occurred in response to unobserved earlier matings, since sucks do not only occur immediately after mating (Schärer et al. 2004).

The significant evolutionary correlation between the presence of reciprocal mating and the purely morphologically-derived reciprocal inferred mating syndrome (Figure 5b) confirms previous findings (Schärer *et al*., 2011). It shows that persistence and resistance are not generally limited to single traits, but are often composite suites of behavioural, morphological and physiological traits acting together (Arnqvist & Rowe, 2005). For example, the thickened female antrum wall and the suck behaviour might be different components of female resistance. While the former might prevent injury resulting from the male genitalia when mating reciprocally, the suck behaviour serves to remove unwanted ejaculate received during mating. Similar adaptations of the female reproductive tract are also seen in the seed beetle *C. maculatus*, where a thicker female tract lining serves as a resistance trait against harm by male genitalia (Dougherty *et al*., 2017). Moreover, resistance and persistence traits can also occur at the proteomic level. A study in *M. lignano* identified two seminal fluid transcripts, experimental knock-down of which caused mating partners to suck more often (Patlar *et al*., 2020). This suggests that the seminal fluid proteins derived from these transcripts might be counter adaptations by the donor to prevent the suck behaviour by the recipient.

In our dataset, there was one species each that exhibited the hypodermic inferred mating syndrome morphology and showed both reciprocal mating and the suck behaviour (*M. rostratum*), only reciprocal mating (*M. distinguendum*), or only the suck behaviour (*M. finlandense*) (Supplementary Table S1). Interestingly, the three species represent at least two, but possibly three, of the above-mentioned multiple independent origins of the hypodermic inferred mating syndrome (Brand *et al*., 2022b). Conversely, there were 12 species that exhibited the reciprocal inferred mating syndrome, but in which neither reciprocal mating, nor the suck behaviour was observed. As above, this mismatch occurred mainly in species for which we had relatively few observation hours (Supplementary Figure S3), suggesting that, if these species have a low mating frequency, then more observation time may be needed to avoid falsely inferring the absence of the mating and suck behaviour. And finally, for many of the species that showed the hypodermic inferred mating syndrome we had considerable amounts of observation hours (Supplementary Figure S3), so that it seems unlikely that the absence of mating and suck observations in these species were due to a lack of effort.

*Macrostomum* species showed large interspecific variation in behaviour, with a nearly 900-fold variation in mating duration, a 3-fold variation in the suck duration, and a nearly 400-fold variation in the mating and suck frequency across the genus (Figure 6). Remarkably, despite this extensive interspecific variation in behavioural traits, we see clear correlations between both the mating and suck duration, as well as the mating and suck frequency, suggesting that the mating and suck behaviour have coevolved. If a longer mating duration or more frequent mating implies more sperm transfer, then we expect selection for a longer suck duration and/or a more frequent suck behaviour (particularly if ejaculate receipt is associated with fitness costs). In some species, at least, a longer mating duration does imply more ejaculate transferred (Engqvist & Sauer, 2003), and is often used as a proxy for ejaculate size (Kelly & Jennions, 2011). Alternatively, such a correlation could also emerge as a result of variation in genital complexity, e.g. if it takes longer to insert and remove more complex male genitalia, and to suck out ejaculate from more complex female antra. Interestingly in *Macrostomum*, male and female genital complexity are indeed correlated (Brand *et al*., 2022b). Moreover, a positive correlation between reciprocal mating and suck could also appear, if some species do not do well under our laboratory conditions, leading to an overall low behaviour frequency.

Note, however, that we confirmed that the individuals we used for mating movies were adults with visible testes and ovaries, and we also established the robustness of the observed correlations by excluding species in which mating or suck had only been observed in one replicate (Supplementary Table S5). We did not find any correlations between the frequency and duration for either reciprocal mating or suck. While we might have expected mating duration to trade off with mating frequency, mating duration only made up a relatively small percentage of total tine, potentially posing no trade-off. Similarly, if sucking is not very costly, the suck duration and frequency may not trade-off; and could even be positively correlated, since both help to remove ejaculate.

Finally, mating frequency (and possibly mating duration) could be positively correlated with allocation towards the male function (e.g., testes). Indeed, studies in *Macrostomum* have shown interspecific variation in sex allocation towards the male and female functions, such as testes and ovaries (Singh *et al*., 2020b; Brand *et al*., 2022a; Singh & Schärer, 2021). This interspecific variation could potentially relate to the mating behaviour, as we can expect species that have a longer mating duration or higher mating frequency to have larger testes, if longer and/or more frequent mating implies that more sperm are transferred (Janicke & Schärer, 2009). Mating duration could also correlate with the complexity of genitalia, such that more complex genitalia might require longer mating duration (King *et al*., 2009), and future studies should investigate the correlations between different aspects of reproductive behaviour and reproductive morphology in *Macrostomum*.

## Conclusions

Our study provides direct observational evidence for ejaculate removal during the postcopulatory suck behaviour in the species *M. hamatum*, compelling support for the coevolution between the reciprocal mating and suck behaviour, and detailed information in a phylogenetic context on the occurrence and interspecific variation of the suck behaviour. Moreover, we show that reproductive morphology can be a good proxy to infer the behavioural mating strategy. Taken together our study shows the presence of a postcopulatory female behavioural resistance trait that co-evolves with mating strategy and allows manipulation of received ejaculate in a simultaneously hermaphroditic sexual system. Thus, our study adds to the repertoire of information on traits involved in sexual conflict in *Macrostomum* genus and demonstrates the genus as an excellent model system for understanding sexual antagonistic coevolution by allowing us to examine the evolution of diverse female resistance and male persistence traits, spanning behavioural and morphological traits, simultaneously.

## Acknowledgements

We would like to thank Gudrun Viktorin, Lukas Zimmermann, Jürgen Hottinger, Urs Stiefel and Daniel Lüscher for technical support. The help from many people that have been involved in the collection and maintenance of the worms we have used here is acknowledged in the study that presents the large-scale *Macrostomum* phylogeny, but here we here would like to specifically thank Philipp Kaufmann, Gregor Schulte, Dita Vizoso, and Michaela Zwyer for contributing several of the analysed mating movies. This research was supported by grants 31003A_162543 and 310030_184916 of the Swiss National Science Foundation (SNSF).

